# A Savory-based Formulation for Sustainable Management of Early Blight caused by *Alternaria solani* and Preservation of Tomato Fruit Quality

**DOI:** 10.64898/2026.01.20.700539

**Authors:** Farzaneh Lak, Azad Omrani, Mohammad Javan Nikkhah, Amir Mirzadi Gohari, Mogens Nicolaisen, Morteza Abuali, Masoud Ahmadzadeh

## Abstract

Environmental degradation caused by excessive chemical inputs is a major global challenge, highlighting the urgent need for sustainable agricultural solutions. Early blight of tomato (*Solanum lycopersicum* L.), caused by *Alternaria solani*, is a widespread disease with limited eco-friendly control options. This study evaluated three savory-based formulations; CC2020, CCP40, and CCE40 each containing savory essential oil with different compositions for managing early blight under both in vitro and greenhouse conditions. Among them, CC2020 demonstrated the highest efficacy in reducing disease severity and showed consistent performance under greenhouse conditions. Additionally, CC2020 helped preserve vitamin C content in tomato fruits and reduced melanin concentration in *A. solani*, suggesting a potential dual role in disease suppression and fruit quality maintenance. These findings suggest that CC2020 could serve as a promising, biocompatible formulation for sustainable management of tomato early blight.

## Introduction

The extensive use of synthetic pesticides has long raised global concerns due to their contribution to environmental pollution, ecological imbalance, and risks to human health. Since the mid-20th century, researchers have repeatedly warned about the persistence and bioaccumulation of these chemicals in natural ecosystems. Recent studies further indicate that conventional pesticides continue to contaminate soil, water, and non-target organisms, leading to long-term ecological and toxicological consequences (Nehul et al., 2025; Rasool et al., 2023; Sharma et al., 2022). These challenges have intensified the demand for environmentally sustainable alternatives, particularly plant-based formulations that offer safer and more degradable options for disease management.

Tomato early blight is one of the most significant and widespread diseases affecting tomatoes (Hou et al., 2021; Khan and Narvekar, 2020). Severity of infection has been reported to range from 19.2% to 67.5%, depending on study conditions and outbreak severity (Perveen and Bokahri, 2020; Adhikari et al., 2017; Datar and Mayee, 1981). Environmental factors, particularly temperature and relative humidity, play a crucial role in disease progression and the extent of crop damage (Attri et al., 2024).

The optimal temperature range for the spread of tomato early blight is 22–30°C, with humidity levels between 74.1% and 83.7% and an average rainfall of 47.07 mm (Fagodiya et al., 2022). Initial symptoms appear as dark brown to black spots on leaves that enlarge and develop concentric rings. As the disease progresses and spores are abundantly produced, symptoms may spread to stems, branches, leaves, and fruits. Under favorable environmental conditions, the causative fungus can cause leaf drop, branch drying, and premature fruit fall, resulting in yield losses of up to 79% (Adhikari et al., 2017; Tanvir et al., 2020).

Most tomato cultivars are susceptible to early blight (Zhang et al., 2003), and limited information is available regarding non-host resistance of tomato cultivars to this necrotrophic, toxin-producing pathogen (Gulzar et al., 2021). The two main causative species have different temperature requirements: *A. solani* thrives at lower temperatures, whereas *A. alternata* requires higher temperatures for disease development (Stammler et al., 2013). Although chemical treatments can manage *Alternaria* infections in tomatoes, prolonged use may lead to resistance in the pathogenic fungus. More importantly, these chemicals contribute to environmental pollution and pose toxicological risks to humans and non-target organisms (Ahmad et al., 2022).

Current efforts to control tomato early blight primarily focus on biological control and the use of biocompatible materials (Ahmad et al., 2022). To date, no fungicide has been identified that is both highly effective and environmentally safe against this pathogen (Alizadeh-Moghaddam et al., 2024). In addition, herbal materials have also demonstrated considerable efficacy in disease management (Kagale et al., 2004).

One environmentally friendly strategy for managing tomato early blight is the use of plant essential oils. Interest in replacing synthetic chemicals with these natural compounds has been increasing, mainly due to concerns about chemical contamination of natural ecosystems. While practices such as crop rotation, use of certified seeds, and maintaining hygiene at production sites can aid in disease management, chemical treatments are often still required for more comprehensive control. Nevertheless, essential oils represent a promising alternative (Mona et al., 2016).

Essential oils, such as thyme (*Thymus vulgaris*), have attracted increasing attention for their strong antifungal properties. Recent studies demonstrate that these natural compounds can effectively inhibit *A. solani* growth and suppress early blight development, offering a sustainable and environmentally friendly alternative to chemical fungicides (Smith and Kumar, 2022).

A recent study (Lurwanu and Muhd, 2025) developed a combination of plant extracts, including neem (*Azadirachta indica*), garlic (*Allium sativum*), and ginger (*Zingiber officinale*), to enhance their stability and efficacy against *A. solani*. The results demonstrated a significant reduction in disease severity under field conditions, while maintaining the physical and chemical quality of tomato fruits. Compared to conventional chemical fungicides, this formulation posed lower environmental risks and was consistent with sustainable agricultural practices.

Another study investigated the synergistic effects of thyme extract in combination with other plant extracts, including garlic (*Allium sativum*) and neem (*Azadirachta indica*), for controlling early blight in tomatoes. The results demonstrated that these combinations exerted enhanced inhibitory effects on *A. solani* growth and significantly reduced disease incidence, while preserving fruit quality. Such plant-based formulations have the potential to decrease the use of chemical fungicides, thereby promoting environmental sustainability and consumer safety (Lee & Patel, 2024).

Many studies have demonstrated the ability of essential oils to inhibit the growth of pathogenic fungi, with some exhibiting fungicidal properties and others showing fungistatic effects. The effectiveness of essential oils in either killing or inhibiting fungal growth depends on their active compounds (Omar and Kordali, 2019).

In this study, plant-based formulations derived from savory (*Satureja* spp.) were evaluated against *A. solani*, the causative agent of early blight in tomatoes, both in vitro and in vivo. The formulations were developed using savory extracts with documented antifungal properties, selected based on traditional knowledge and recent scientific findings. The goal was to maximize efficacy through synergistic interactions among the components.

Beyond their antifungal activity, the savory-based formulations also preserved key nutritional qualities of tomato fruits, such as vitamin C content, indicating a dual role in both disease management and postharvest quality maintenance. This eco-friendly plant-based treatment provides a sustainable alternative to synthetic fungicides, with additional benefits for food quality and consumer health. Vitamin C is a key water-soluble bioactive compound in tomatoes (*Solanum lycopersicum*) (Prabha et al., 2011). It plays multiple roles in plants, contributing to both overall health and nutritional value by helping plants cope with various stresses and enhancing the production of certain plant hormones (Sharma et al., 2024). Tomatoes are valued globally not only for their yield but also for their nutritional attributes, particularly vitamin C (ascorbic acid), which functions as a potent antioxidant and a critical quality marker for consumer acceptance (Raiola et al., 2014; Lucier et al., 2000). Vitamin C content in fruits can vary depending on cultivar, ripening stage, and environmental or management conditions (Martí et al., 2018). Among biotic stresses, pathogen attacks can influence the ascorbate pool in plant tissues, often leading to oxidative stress or altered antioxidant metabolism (Noctor & Foyer, 1998). Therefore, evaluating the effects of biological control strategies on both disease suppression and nutritional outcomes, such as vitamin C content, can enhance the multifunctionality of plant-based products (González et al., 2021). Vitamin deficiencies in processed foods can lead to various health disorders and syndromes in humans (Sebrell et al., 1969). The use of natural materials and organic inputs in crop cultivation influences the concentration of essential nutrients, including vitamin C (Crinnion et al., 2010). Additionally, treatment of agricultural products with bio-based inputs has been shown to increase their vitamin content (Williams et al., 2002)

Melanin is widely distributed in fungi and plays a key role in enhancing tolerance to environmental stresses, particularly ultraviolet radiation, enzymatic degradation, and high temperatures (Cho et al., 2012). While melanin production is not strictly essential for pathogenic fungi, it reinforces protective mechanisms against external stressors (Zeng et al., 2020; Krishnan et al., 2018). However, the absence of melanin can reduce the fungus’s ability to penetrate host tissues (Wang et al., 2023). Melanin is a dark pigment produced by several phytopathogenic fungi, including *A. solani*, and plays a key role in fungal pathogenicity and survival. It protects fungal cells from environmental stresses such as UV radiation, oxidative bursts from host plants, and antifungal compounds (Bell and Wheeler, 1986). In *Alternaria* species, melanin accumulation in cell walls enhances resistance to plant defense mechanisms, increasing virulence and persistence in host tissues (Langfelder et al., 2003; El-Banna et al., 2018). Therefore, monitoring changes in melanin concentration can provide valuable insights into how fungal virulence is influenced by biocompatible treatments, such as CC2020.

### Laboratory tests

#### Cultivation of fungal isolates

The purified isolates were examined for morphological characteristics, and the fungal species were identified according to the criteria described in *Identification of Alternaria* Species (Simmons, 2007). Additionally, all isolates were processed by the Plant Diseases Department of the National Research Institute in Iran. For purification, *Alternaria* isolates were subjected to monosporulation on Potato Dextrose Agar (PDA) and incubated at 25°C for 3–5 days until growth was established. The isolates were then subcultured to obtain pure cultures.

#### Plant material

The tomato cultivar 4129 used in this study was obtained from Department of Greenhouse Research, Tehran Agricultural and Natural Resources Research and Training Center, Agricultural Research, Education and Extension Organization (AREEO), Tehran, Iran. This cultivar widely used by greenhouse growers in Iran,

#### Experimental Sites

This study was conducted at multiple locations: the Biological Laboratory at Tehran University, the Pathology Laboratory at Middle East Fruit Science Company, the Agricultural and Natural Resources Research Center of Tehran Province (research greenhouse), and a tomato net house in Karaj, Iran.

#### Plant-based formulations

The compounds were prepared using essential oils supplied by the Middle East Fruit Science Company. Among the essential oils of rosemary, savory, mint, thyme, and lemongrass, savory (*Satureja* spp.) essential oil was selected for formulation due to its strong antifungal activity against *A. solani*, as confirmed by preliminary antifungal assays and supported by previous studies. The formulations were chosen based on this screening and their potential for practical application in crop protection. The chemical composition of the savory essential oil used in the formulations was provided by the supplier and is summarized in Table 1. GC-MS analysis was conducted at the Medicinal Plants Research Center of the Research Institute of Forests and Rangelands, affiliated with the Forests, Range, and Watershed Management Organization in Tehran Province, Iran.

**Table 1.** Chemical composition of *Satureja* spp. essential oil.

| Entry | Retention Time (min) | Compound | Area % (GC) | Retention index (RI) |
| --- | --- | --- | --- | --- |
| 1 | 8.27 | α-thujene | 0.41 | 930 |
| 2 | 8.57 | α-pinene | 0.36 | 939 |
| 3 | 10.35 | myrcene | 0.85 | 988 |
| 4 | 11.86 | *p*-cymene | 3.76 | 1031 |
| 5 | 13.17 | γ-terpinene | 0.54 | 1064 |
| 6 | 13.71 | cis-sabinene hydrate | 0.40 | 1077 |
| 7 | 14.82 | linalool | 0.92 | 1101 |
| 8 | 18.57 | Terpinen-4-ol | 0.45 | 1171 |
| 9 | 20.89 | Carvacrol methyl ether | 0.29 | 1243 |
| 10 | 23.24 | thymol | 0.41 | 1295 |
| 11 | 24.02 | carvacrol | 90.32 | 1311 |
| 12 | 32.10 | β-bisabolene | 0.96 | 1508 |
| 13 | 35.17 | Caryophyllene oxide | 0.34 | 1575 |

**Legend:** The table presents the retention time (RT), retention index (RI), and area percentage of each compound in *Satureja* spp. essential oil as determined by GC-MS analysis

#### Determination of the Minimum Inhibitory Concentration (MIC) of Plant-Based Formulations in Vitro

Under laboratory conditions, eight concentrations of each formulation (1000, 1500, 2000, 2500, 3000, 3500, 4000, and 4500 ppm) were prepared and tested to determine the minimum inhibitory concentration (MIC). The medium, PDA (39 grams per liter of water), was prepared, and ceftriaxone was added at 10 mg/L to prevent bacterial contamination. Approximately 20 mL of the prepared medium was dispensed into each Petri dish for the assay.

The formulations were incorporated into the PDA medium at the specified concentrations and sterilized by autoclaving at 121°C for 15 minutes. Once the culture medium cooled to an appropriate temperature and solidified, a 5 mm diameter fungal disk was extracted from a 7-day-old fungal culture and placed at the center of the treatment and untreated Petri dishes.

All Petri dishes were incubated at 25°C. After seven days, the diameter of fungal colony growth in both treated and untreated dishes was measured to evaluate the effectiveness of the plant-based formulations. The lowest concentration that completely inhibited the growth of *A. solani* was recorded as the minimum inhibitory concentration (MIC) (Plodpai et al., 2013).

#### Evaluation of the effect of studied compounds on the mycelium morphology of *A. solani*

To assess the morphological changes induced by the plant-based formulations, *A. solani* was cultured on PDA medium supplemented with different concentrations below the minimum inhibitory concentration (sub-MIC). Fungal plugs (6 mm in diameter) were placed at the center of the plates and incubated at 25 ± 1°C for five days in the dark. After incubation, the mycelial growth was collected and examined under a light microscope at ×400 magnification to observe alterations in hyphal structure, branching, pigmentation, and cytoplasmic integrity compared to the untreated control (Li et al., 2015).

#### The effect of different treatments on melanin concentration Fungal growth for identifying accumulated pigments

Eight-millimeter mycelial blocks were taken from the margin of a 7-day-old *A. solani* culture previously grown on PDA medium using a sterile cork borer. The blocks were placed at the center of Petri dishes containing PDA medium supplemented with different sub-MIC concentrations of the formulations. After seven days of incubation, the fungal culture was soaked in sterile distilled water, scraped with a sterile scalpel, and passed through a metal strainer to remove excess water. The collected material was transferred into sterilized Falcon tubes and stored at −20°C until melanin extraction (modified from Fernandes et al., 2015).

#### Extraction of melanin from fungi

To extract melanin from the fungal mycelium, the dried mycelium was ground using a mortar, and 1 M NaOH was added. The mixture was then autoclaved at 121°C for 30 minutes. After centrifugation at 4000 rpm for 10 minutes, the pigment was recovered from the supernatant (Rajagopal et al., 2011).

#### Quantification of melanin extracted from fungus

To quantify the fungal pigment, the concentration of the pigment obtained from the previous steps was measured using spectrophotometry at a wavelength of 340 nm.

### Testing plant-based formulations in greenhouse conditions

#### Pathogenicity test of *A. solani*

The pathogenicity test for these isolates was conducted in vitro under optimal temperature and humidity conditions for the pathogenic fungus. Three pots were inoculated with a spore suspension at a concentration of 1 × 10^6^ spores/mL for each fungal isolate, while one untreated pot per treatment was inoculated with sterile distilled water without fungal spores (Kamalakhan et al., 2008).

#### Tomato cultivation in a greenhouse: Inoculation with *A. solani* and treatment with plant-based formulations and the synthetic fungicide Chlorothalonil

Tomato seeds were surface-sterilized using a 1% sodium hypochlorite solution for one minute, followed by three successive rinses with sterile distilled water. After drying on sterile filter paper, the seeds were sown in specialized seedling trays. At the two-leaf stage, seedlings were transplanted into pots containing a cocopeat and perlite growing medium mixed in a 3:2 ratio. Tomato plants at the three-to five-leaf stage were used for each treatment (Nashwa and Abo-Elyousr, 2012).

To prepare the spore suspensions of *A. solani* isolates, fresh seven-day-old cultures were used. Seven-week-old tomato plants were then selected for inoculation under controlled greenhouse conditions, with 60% humidity and a 14-hour photoperiod. The average daytime and nighttime temperatures were maintained at 25°C and 20°C, respectively.

In this study, concentrations of 4000, 6000, and 8000 ppm of each selected plant-based formulation were chosen based on preliminary screening of their efficacy against *A. solani*. The formulations were sprayed onto the plants using 8-mL laboratory spray bottles, each delivering approximately 50 μL per puff, with three puffs applied to each leaf to ensure uniform coverage.

To assess the curative effect, plants were first inoculated with spore suspensions at a concentration of 1 × 10^6^ spores/mL and then treated with the plant-based formulations 24 hours later (Stammler et al. 2014). This treatment was repeated three times at five-day intervals.

The same protocol was applied for a chemical fungicide (Chlorothalonil) as a positive control. Disease symptoms and progression were assessed 15 days after inoculation (Bajpai and Kang, 2010). A control treatment was included for each compound, in which plants were inoculated with sterile distilled water without fungal spores.

#### Assessment of Disease Severity and Disease Index under Greenhouse Conditions

The severity of early blight symptoms on tomato leaves was visually estimated for each treatment using a disease rating scale ranging from 0 to 4, as described below. Disease severity was assessed based on the percentage of symptomatic leaf area. A score of 0 was assigned to healthy plants with no visible symptoms; 1 for 1–25% infection; 2 for 26–50% infection; 3 for 51– 75% infection; and 4 for 76–100% infection of on the leaf surface. The Disease Index (DI) under greenhouse conditions was calculated using the following formula described by Hubballi et al. (2010):

***Disease Index (DI) = [Σ (Disease rating × Number of plants in that rating)] / (Total number of plants × Maximum rating) × 100***

#### Preservation of ascorbic acid content in tomato fruits following plant-based treatments Vitamin C assessment

Tomato fruits were sprayed three times at five-day intervals prior to harvest with the formulation at concentrations of 4000, 6000, and 8000 ppm, following the treatment regime used for managing early blight disease. For each treatment, three plants were selected as replicates. Following harvest, fruits were stored at 13 °C, and vitamin C content was evaluated at 0, 5, 10, and 15 days of storage (modified from Nour et al., 2015).

A total of 100 grams of the selected fruit was placed into a model blender and finely ground. During grinding, 50 mL of distilled water was added to the mixture. After mixing, the resulting puree was passed through a strainer to obtain a uniform consistency. The final sample was then transferred into 100 mL flasks (modified from Stpathy et al., 2021).

A 1% (w/v) starch solution was prepared by dissolving 0.5 g of starch in 50 mL of distilled water heated to near boiling. The mixture was stirred continuously until the starch completely dissolved and then cooled to room temperature before use. Subsequently, 1.5 g of potassium iodide (KI) and 0.268 g of potassium iodate (KIO_3_) were dissolved in 200 mL of distilled water with continuous stirring until fully dissolved. The titration was then carried out according to the following steps:

Add 20 mL of the prepared sample in a 50 mL erlen, then 10 mL of starch solution was added to the sample container as a reagent. The prepared solution is poured into the burette and the initial volume is recorded. The container containing the prepared sample is placed under the broth and little by little iodine solution is added to it until it reaches the desired point, and that is when the first color change of the sample turns blue, and at this time, the contents in the sample container are stirred for 20 seconds until the color change is fixed and at this point the addition of iodine solution stops and the secondary volume of iodine solution is read and noted in the burette. This step should be repeated at least twice. Finally measurement is done.

#### Statistical analysis

All experiments were conducted in triplicate with independent replicates to ensure data repeatability. Disease incidence and fruit quality parameters were recorded for each replicate. Data were analyzed using analysis of variance (ANOVA) in SAS software (version 9.4). Treatment means were compared using Duncan’s multiple range test at a significance level of *p* < 0.05. Error bars in figures represent standard deviation (SD) calculated from the three independent replicates. The chemical fungicide (Chlorothalonil) was used as a positive control only in greenhouse experiments, allowing a direct comparison of the effects of plant-based formulations under controlled conditions. This approach provides transparency in the analytical procedures and allows clear assessment of the effects of plant-based formulations and controls on the measured parameters.

## Results

### Screening of plant-based formulations in vitro

The results obtained were remarkable and suggest a promising future for this generation of eco-friendly compounds. These compounds effectively inhibited fungal growth and exhibited both fungicidal and fungistatic properties (Figure 1). Among the tested formulations, the CC2020 compound demonstrated the best performance, followed by CCP40 and CCE40. However, there was no significant difference between the latter two.

**Figure 1.**
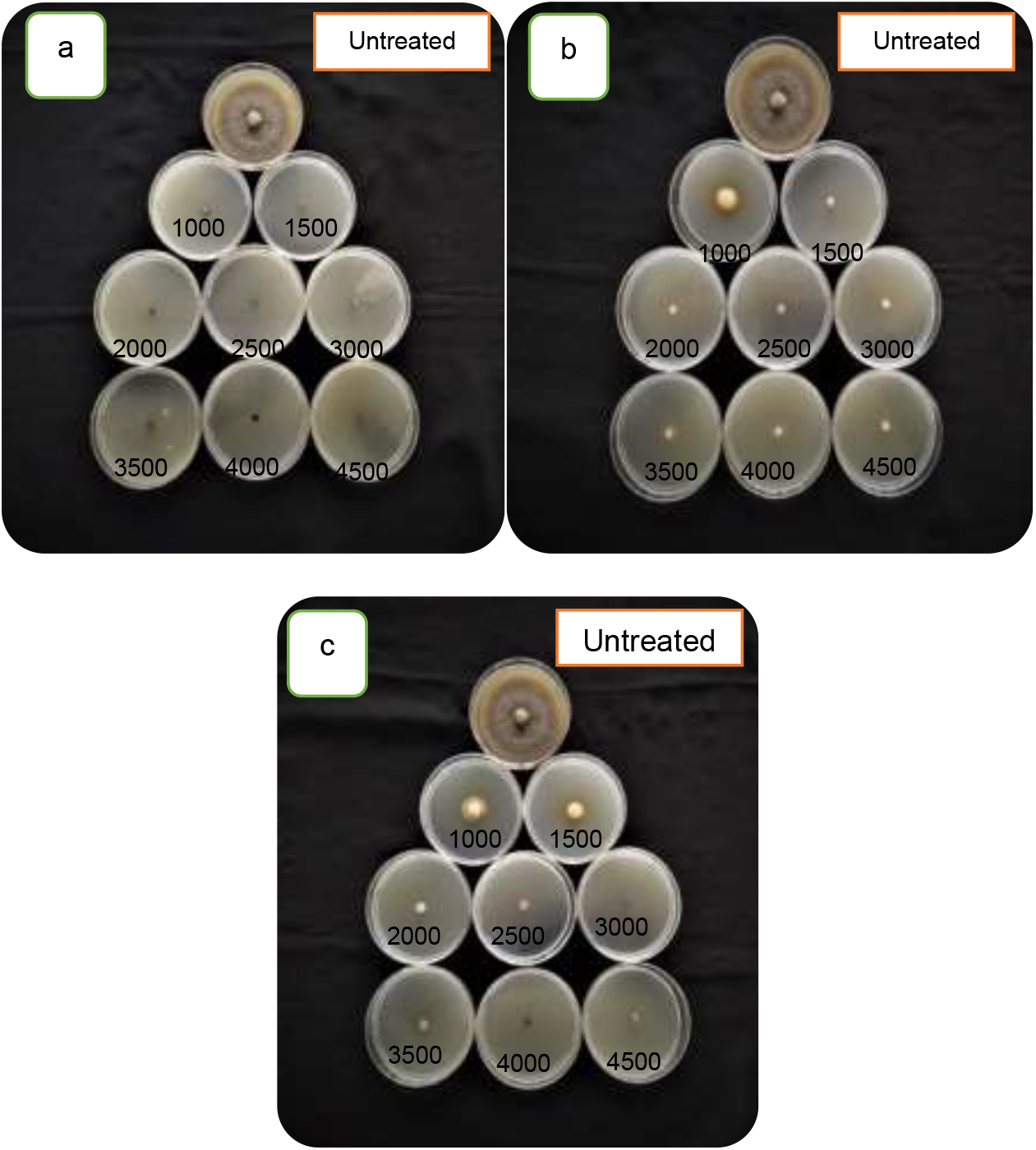
Effect of CC2020 (a), CCP40 (b), and CCE40 (c) compounds on fungal mycelial inhibition at concentrations of 1000, 1500, 2000, 2500, 3000, 3500, 4000, and 4500 ppm. The minimum inhibitory concentrations demonstrating significant antifungal effects for each compound are indicated

### MIC determination of the biocompatible formulations

The minimum inhibitory concentrations (MICs) of the different formulations are presented in Table 2. Among the tested formulations, CC2020 exhibited the lowest MIC value (1000 ppm), CCP40 showed an intermediate MIC (1500 ppm), and CCE40 had the highest MIC (2000 ppm).

**Table 2.** Minimum inhibitory concentrations of different plant-based formulations *Alternaria solani*.

| Treatment | MIC (ppm) |
| --- | --- |
| CC2020 | 1000 |
| CCP40 | 1500 |
| CCE40 | 2000 |

**Legend:** This table shows the minimum inhibitory concentrations (MIC) of three different plant-based formulations tested against the fungal pathogen *A.solani*. MIC values are expressed in parts per million (ppm).

### Assessment of the impact of the studied compounds on the mycelial morphology of *A. solani*

The study revealed that the mycelium of *A. solani* was markedly affected by the plant-based formulations. All treatments induced distinct morphological changes, including plasmolysis (separation of the cytoplasm from the cell wall), loss of cytoplasmic contents, cellular degeneration, and eventual cytoplasmic disintegration. Microscopic observations supported these findings: treated samples showed severely degraded or absent cellular contents, whereas the untreated control maintained intact and clearly visible internal structures. Although the antifungal effects were generally similar among the formulations, CC2020 completely eliminated fungal cells and hyphal structures, highlighting its particularly potent activity (Figure 2).

**Figure 2.**
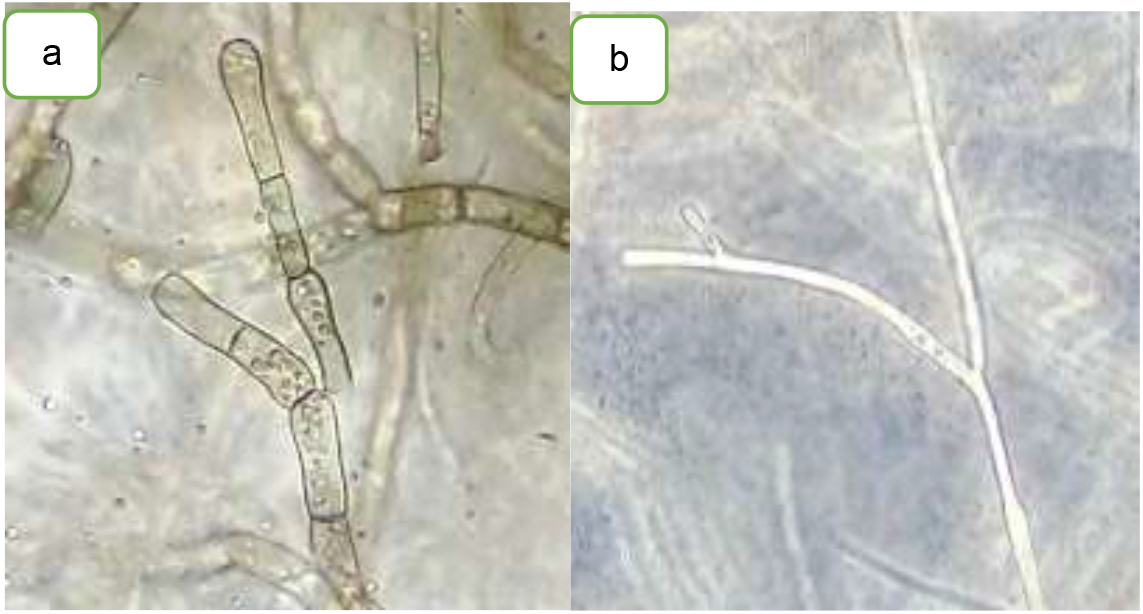
Untreated mycelium (control, a) shows intact cytoplasmic content, whereas mycelium treated with plant-based formulations CC2020, CCP40, and CCE40 (b) exhibits cytoplasmic degradation. All three formulations induced similar effects, and a representative image is shown for the treated samples.

### The Effect of different treatments on melanin levels

To assess the effect of plant-based formulations on melanin production by *A. solani*, melanin was extracted and quantitatively measured using a spectrophotometer. Statistical analysis was performed on the measured values, and the results are presented in Figure 3.

**Figure 3.**
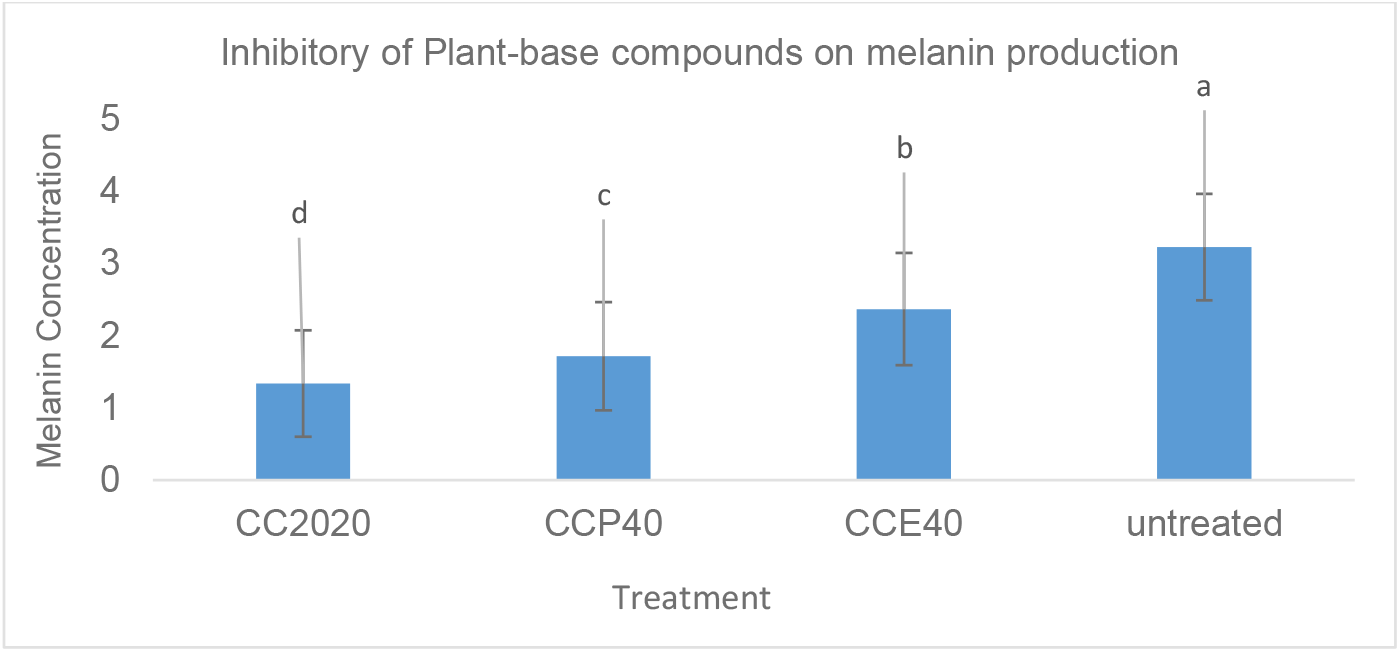
Effect of plant-based formulations on melanin production in *A. solani*. Error bars represent standard deviation from three independent replicates. Statistical analysis was performed using SAS software, and treatment means were compared using Duncan’s multiple range test (p < 0.05). Different letters (a–d) indicate statistically significant differences.

### Result related to greenhouse examinations

#### Pathogenicity test

The *A. solani* isolate was tested for pathogenicity on the tomato. The results confirmed that *A. solani* is pathogenic to tomatoes. The incidence of disease (DI) of pathogenicity for both the isolates and the untreated are presented in Table 3.

**Table 3.** Comparison of Disease Index (DI)

| Treatments | Disease Index |
| --- | --- |
| *A.solani* | 83.33% |
| Untreated | 0% |

**Legend:** This table compares the Disease Index (DI) among different treatments. Values are expressed as percentages.

#### Evaluation of the effect of plant formulations on early blight disease symptoms under greenhouse conditions

According to Table 4, the highest mean disease incidence was observed in the control treatment (distilled water), while the lowest occurred in the treatments with plant-based formulations.

**Table 4.** Comparison of the Disease Index (%) among treatments.

| Treatments | Dose | Disease Index |
| --- | --- | --- |
| CC2020 | 4000 ppm | 25% |
| CC2020 | 6000 ppm | 8.33% |
| CC2020 | 8000 ppm | 8.33% |
| CCP40 | 4000 ppm | 25% |
| CCP40 | 6000 ppm | 16.66% |
| CCP40 | 8000 ppm | 8.33% |
| CCE40 | 4000 ppm | 25% |
| CCE40 | 6000 ppm | 25% |
| CCE40 | 8000 ppm | 16.66% |
| Clorotalonil | 2000 ppm | 8.3% |
| untreated | - | 100% |

**Legend:** This table presents the curative effects of different treatments against early blight in tomato, expressed as Disease Index (DI) percentages.

Greenhouse studies were conducted to evaluate the curative effects of plant formulations and the chemical fungicide Chlorothalonil.

The effectiveness of different plant-based formulations (CC2020, CCP40, and CCE40) at three concentrations (4000, 6000, and 8000 ppm), as well as the chemical fungicide Chlorothalonil at the recommended dose of 2000 ppm, was evaluated under greenhouse conditions.

To assess their curative effects, a spore suspension of *A. solani* (1×10^6^ spores/mL) was prepared and sprayed onto the plants. After 24 hours, the plant formulations or Chlorothalonil were applied. Disease symptoms developed on untreated control plants sprayed with sterile distilled water only, whereas plants treated with the formulations or Chlorothalonil exhibited markedly fewer symptoms. The infected control group showed a higher disease incidence compared to the treated plants, demonstrating the effectiveness of the plant-based formulations in reducing early blight under greenhouse conditions. The results are presented in Figure 4.

**Figure 4.**
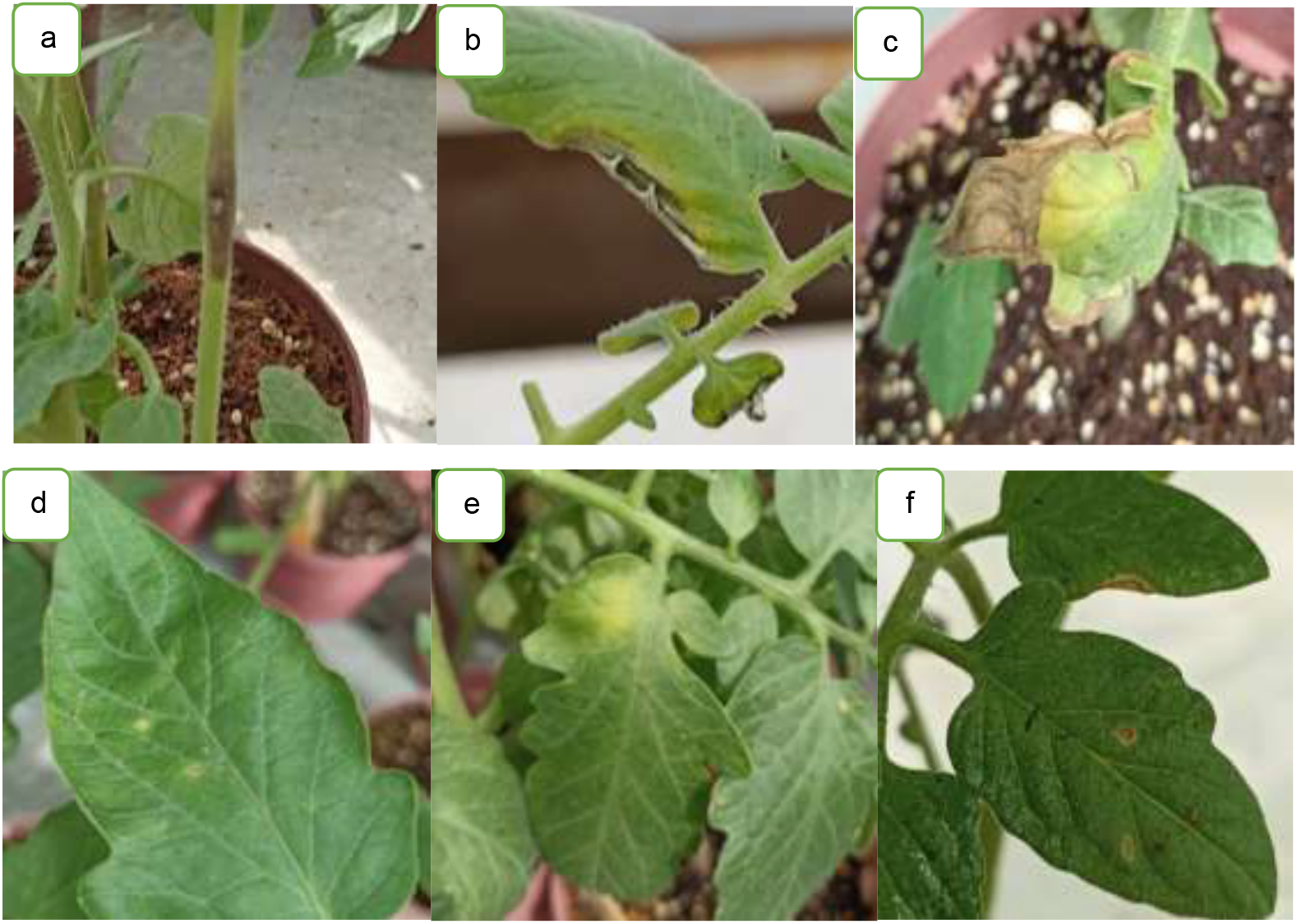
Panels (a), (b), and (c) show the untreated control plants exhibiting the highest disease severity, while panels (d), (e), and (f) illustrate the effects of the plant-based formulations CC2020, CCP40, and CCE40, respectively, in reducing early blight symptoms in tomato under greenhouse conditions.

The results indicated a significant difference between the treatments applied for controlling early blight symptoms on cultivar 4129 as illustrated in Figure 5. According to this bar chart plant based and chemical compounds are able to control this disease efficiency compared to water treatment.

**Figure 5.**
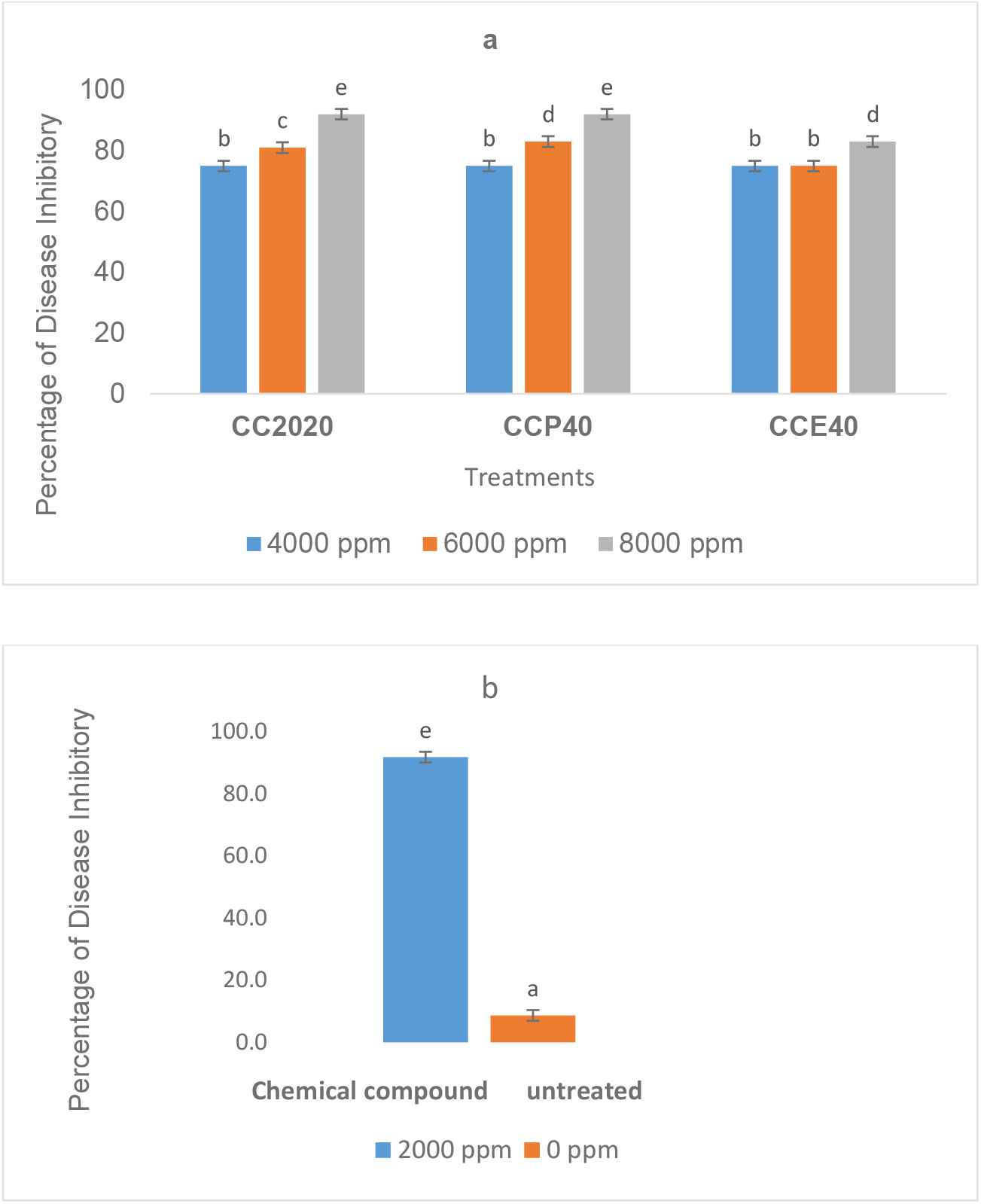
Effectiveness of plant-based formulations at concentrations of 4000, 6000, and 8000 ppm (panel a), and of the chemical fungicide Chlorothalonil (2000 ppm) and water control (0 ppm) (panel b) in controlling early blight in tomato. Statistical analysis was performed using SAS software, and treatment means were compared using Duncan’s multiple range test at p < 0.05. Different letters (a–d) indicate statistically significant differences.

#### Evaluation of vitamin C

The effects of plant-based formulations on vitamin C content in tomato fruits were evaluated at three storage times: day 0, 10, and 15 post-harvest at 15 °C (Figure 6).

**Figure 6.**
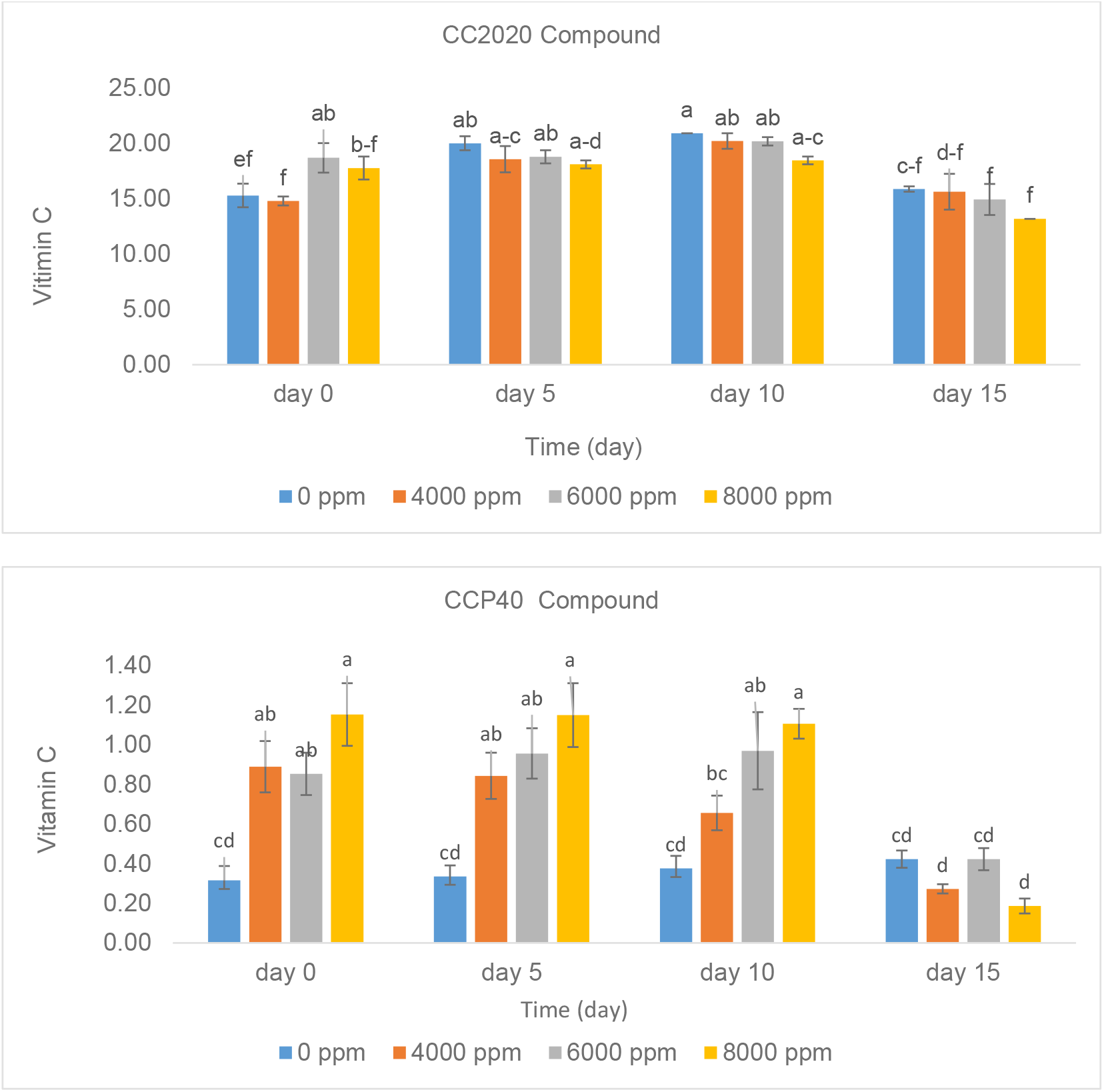
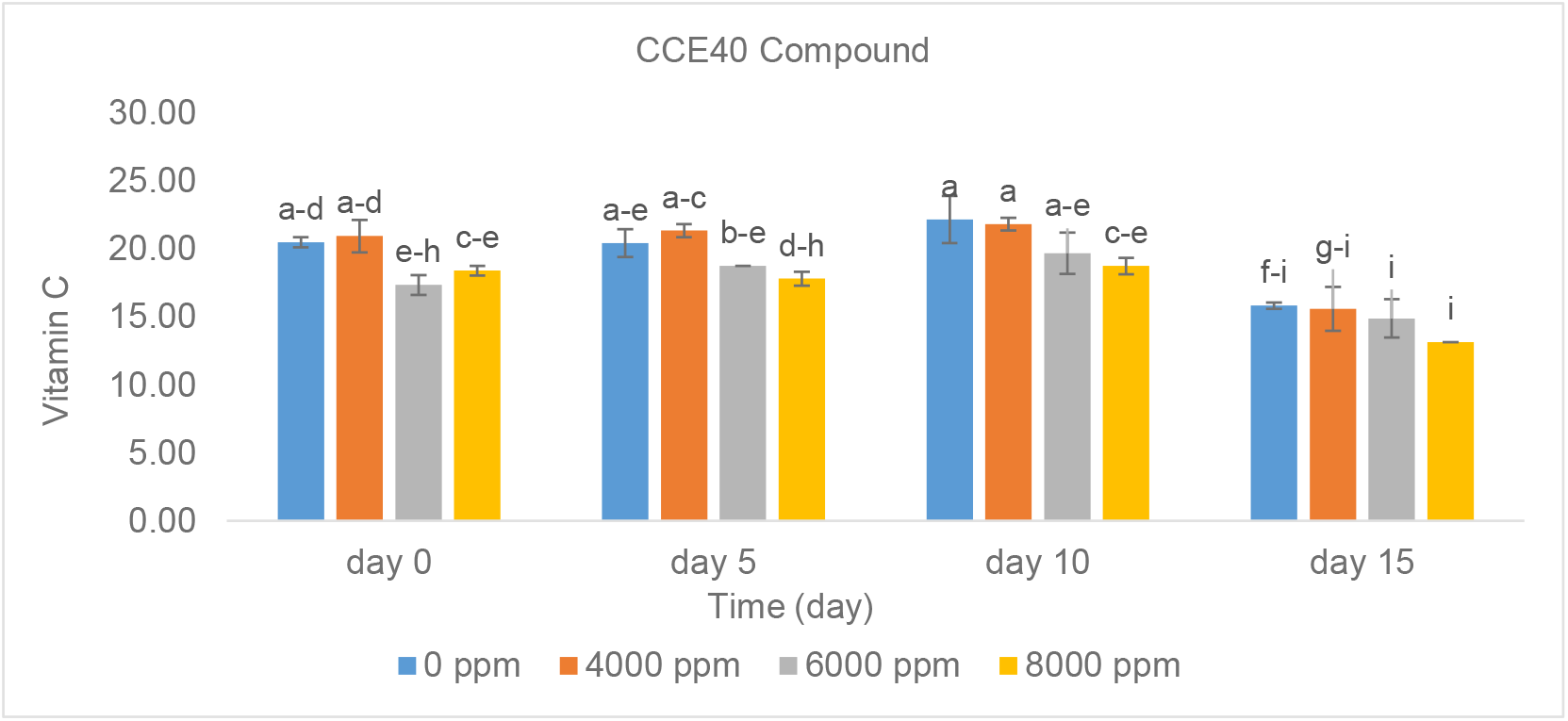
Effect of plant-based formulations applied before harvest at concentrations of 4000, 6000, and 8000 ppm on the vitamin C content of tomatoes. Statistical analysis was performed using SAS software, and treatment means were compared using Duncan’s multiple range test at *p* < 0.05. Different letters (a–d) indicate statistically significant differences. Error bars represent standard deviation from three independent replicates

#### Effect of CC2020 on Vitamin C

CC2020 at 6000 ppm maintained the highest vitamin C levels across all storage days, indicating optimal preservation. In contrast, 8000 ppm showed a decline over time, suggesting that higher concentrations may negatively affect retention. Vitamin C content decreased in all treatments over the storage period, with the untreated control showing the lowest levels by day 15. Differences in letter groupings indicate that the effect of CC2020 is influenced by storage duration.

#### Effect of CCP40 on Vitamin C

CCP40 at moderate concentrations (4000–6000 ppm) effectively preserved vitamin C levels over the storage period. In contrast, 8000 ppm resulted in significantly lower vitamin C content, suggesting that higher doses may negatively affect retention. Vitamin C levels generally decreased over time in all treatments, with the untreated control showing the lowest levels by day 15. These results indicate that moderate CCP40 concentrations provide optimal protection for tomato vitamin C during storage.

#### Effect of CCE40 on Vitamin C

CCE40 maintained vitamin C levels most effectively at moderate concentrations (4000– 6000 ppm). In contrast, the 8000-ppm treatment resulted in a marked reduction in vitamin C content, suggesting that higher doses may induce stress or accelerate degradation. Among the tested concentrations, 6000 ppm was statistically the most effective for preserving vitamin C during storage, whereas 8000 ppm was the least effective.

## Discussion

Previous studies on the management of early blight have predominantly focused on the antifungal activity of individual essential oils against *A. solani*. In contrast, the present study evaluated three distinct plant-based formulations and demonstrated that all of them possess substantial antifungal potential. Among these, the formulation CC2020 consistently exhibited the highest level of efficacy across the assays. The overall efficacy of these formulations was comparable to that of the chemical fungicide Chlorothalonil, challenging the prevailing assumption that plant-based compounds are inherently less effective than synthetic fungicides.

Given their natural origin, absence of pre-harvest interval (PHI) restrictions, and comparatively lower environmental and health risks, these formulations may be considered promising candidates for sustainable disease management. To the best of our knowledge, there are limited reports evaluating plant-based formulations with comparable structural features that exhibit such broad and consistent antifungal activity against *A. solani*.

Several studies have documented the antifungal activity of savory (*Satureja* spp.) essential oils against phytopathogenic fungi, including *A. solani*. The major constituents of savory oil, particularly carvacrol and thymol, are widely recognized for their strong antimicrobial properties. Sharifi-Rad et al. (2015) reported substantial inhibition of *A. solani* mycelial growth following exposure to *Satureja hortensis* essential oil under in vitro conditions. Likewise, Mohammadi et al. (2018) demonstrated that savory oil effectively reduced spore germination and decreased lesion development on tomato leaves. These findings strongly support the inclusion of savory essential oil as a key active component in the plant-based formulations evaluated in the present study.

Similarly, Sesan et al. (2016) demonstrated that *Satureja hortensis* extracts significantly reduced *A. alternata* infection in horticultural crops under both in vitro and in vivo conditions. These findings align well with the present study, where savory essential oil served as the primary active ingredient in all evaluated formulations. Its strong antifungal properties substantially contributed to the superior performance of the most effective formulation, CC2020.

Combined essential oils containing *Satureja hortensis* and other plant-derived oils have also demonstrated synergistic antifungal effects against *Alternaria* species. Gholamnezhad et al. (2018) reported that a blend of *S. hortensis* and *Thymus vulgaris* essential oils significantly reduced *A. solani* growth and sporulation, attributing this enhanced activity to the interactive effects of bioactive components such as thymol, carvacrol, and p-cymene, which disrupt fungal membranes and interfere with melanin biosynthesis. Similarly, Abdollahzadeh et al. (2020) showed that combining savory oil with cinnamon oil increased antifungal efficacy compared to individual applications, emphasizing the potential of multi-component plant-based fungicides. These findings further support the rationale for selecting savory oil as the core ingredient in the formulations tested in this study, given its documented potency and compatibility with the formulation objectives.

The antifungal activity of savory essential oil is largely attributed to its capacity to disrupt fungal cell membranes and inhibit ergosterol biosynthesis, a critical component required for maintaining membrane structure and function (Burt, 2004). Additionally, the presence of multiple bioactive constituents—particularly phenolic monoterpenes—creates synergistic interactions that enhance overall antifungal potency while reducing the risk of resistance development. The outcomes of the present study are consistent with these mechanisms, as the savory-based formulations, especially CC2020, demonstrated strong inhibitory effects on *A. solani* and effectively reduced disease progression. These findings further support the potential of savory-derived, multi-component plant formulations as sustainable alternatives to synthetic fungicides in early blight management.

According to Nascimento et al. (2008) and Chutia et al. (2009), essential oils extracted from pepper and citrus species have shown strong inhibitory activity against *Alternaria* spp. Similarly, Mona et al. (2016) demonstrated that lemongrass, thyme, and sweet basil oils effectively suppressed the mycelial growth of *A. solani*. In line with these findings, Babagoli et al. (2012) reported significant antifungal effects of savory, thyme, and ajwain essential oils when incorporated into culture media. Collectively, these studies highlight the broad potential of aromatic plant oils in managing *Alternaria* infections, supporting the rationale behind employing savory-based formulations in the present study.

These inhibitory effects have been consistently confirmed under in vitro conditions. Savory and thyme essential oils, in particular, have demonstrated strong antifungal activity against *A. solani* (Sedigheh Esmaeil et al., 2019; Smith and Kumar 2024). Similarly, Yadav and Mishra (2018) reported comparable inhibitory effects of savory, thyme, and ajwain essential oils against *A. alternata* in culture media. These findings further corroborate the efficacy of savory-based formulations employed in the current study.

Moreover, Singh et al. (2005) identified menthone, methyl acetate, isomenthone, and limonene as the dominant constituents in the essential oil of *Mentha arvensis*, while Babagoli et al. (2012) reported that the essential oil of *Carum capticum* significantly inhibited the growth of *A. solani*.

The observed reduction in melanin content in *A. solani* following treatment with CC2020 may be associated with a weakening of the pathogen’s defenses and virulence. Melanin protects fungal cells against host-derived oxidative stress and environmental challenges and facilitates tissue colonization (Bell and Wheeler, 1986; Langfelder et al., 2003). Accordingly, a reduction in melanin content caused by CC2020 may contribute to decreased pathogen aggressiveness and enhanced sensitivity to plant defense mechanisms.

Beyond its direct antifungal activity, CC2020 was also found to preserve vitamin C content in tomato fruits, suggesting a potential dual functionality (Patel et al., 2020). As ascorbic acid is an important quality attribute influencing shelf life, consumer preference, and nutritional value, this effect may have both agronomic and commercial relevance (Ilahy et al., 2019). The maintenance of ascorbate levels could be associated with induced resistance and activation of plant antioxidant pathways, potentially triggered by bioactive compounds present in the CC2020 formulation (Boubakri et al., 2013). Such dual-action formulations, combining pathogen suppression with nutritional enhancement, may represent a promising approach for sustainable crop protection (Bhanu et al., 2021).

Overall, the literature indicates that essential oils can exhibit strong antifungal activity, with some acting as fungicidal agents and others as fungistatic inhibitors. Their efficacy largely depends on the specific bioactive compounds present in each oil (Omar and Kordali, 2019). In this context, CC2020, as a plant-based formulation, may offer a safer and more environmentally sustainable alternative to conventional fungicides, highlighting its potential role in integrated and eco-friendly crop protection strategies. Its effectiveness has been demonstrated under multiple experimental conditions, including greenhouse trials, open-field experiments, and post-harvest storage, suggesting its suitability for further development and possible commercialization as a sustainable plant protection product.

## Conclusion

This study presents an eco-friendly approach for managing early blight in tomato using plant-based formulations. The results indicate that CC2020 can effectively reduce disease severity, help preserve fruit vitamin C content, and lower melanin levels in *A. solani*, suggesting a dual effect on both pathogen suppression and fruit quality maintenance. These findings imply that CC2020 could serve as a promising alternative to conventional chemical fungicides, with potential applicability under greenhouse. Further research is ongoing to clarify the precise mechanisms underlying its antifungal and protective effects.

